# Immunological cross-reactions with paramyxovirus nucleoproteins may explain sporadic apparent ebolavirus seropositivity in European populations

**DOI:** 10.1101/2020.03.27.007195

**Authors:** Lisa A. Bishop, Marcell Müllner, Amalie Bjurhult-Kennedy, Robert M. Lauder, Derek Gatherer

## Abstract

Although confirmed outbreaks of Ebola Virus Disease (EVD) have been confined to central and west Africa, seropositivity to Zaire ebolavirus (EBOV) has been reported in other parts of Africa and even in one study from the early 1990s in Germany. The possible reasons for the discrepancy between serological studies and clinical evidence, remain uncertain. Here, we report anonymous serum donors sampled in Lancaster (UK) with seropositivity by ELISA to EBOV nucleoprotein at a frequency of approximately 2%. In one serum donor, we confirm the result using Western blot. This is only the second report of seropositivity for EBOV outside of Africa. Our samples are negative to EBOV glycoprotein, suggesting that the seropositivity is specific to the nucleoprotein and may be due to cross-reaction with antibodies produced by exposure to another virus. To investigate potential candidates for this cross-reacting virus, we perform bioinformatics analysis that suggests that EBOV nucleoprotein has structural similarity to paramyxovirus nucleoproteins at a candidate immunological epitope. Cross-reaction of antibodies against paramyxovirus nucleoproteins, with EBOV nucleoprotein antigens used in serological testing, may be the cause of the rare instances of ebolavirus seropositivity in Europe, and may also be a confounding factor in African serosurveys.

## 1. Introduction

Ebola Virus Disease (EVD) is caused by viruses of the genus *Ebolavirus* (Realm *Riboviria*; Phylum *Negarnaviricota*; sub-phylum *Haploviricotina*; Class *Monjiviricetes*; Order *Mononegavirales*; Family *Filoviridae*), and has been considered a serious emerging disease threat in tropical Africa since the discovery of Zaire ebolavirus (EBOV) in 1976. Although confirmed outbreaks of EVD produced by EBOV have been restricted to central and west Africa, the geographical range over which human populations display seropositivity to EBOV and other filovirus antigens is far larger. The serological data prior to 2016 has been reviewed by Formella & Gatherer ^1^, Nyakarahuka et al ^2^ and Bower & Glynn ^3^ and commented on by Crozier ^4^. Since then, further data has been obtained on seroprevalence in both humans ^5–11^ and animals ^12–15^.

Formella & Gatherer ^1^ suggested three potential explanations for the serological signal outside of confirmed outbreak areas: 1) previous full outbreaks of EVD misclassified as other diseases, 2) previous infectious chains of EBOV with a milder, or even asymptomatic, clinical presentation, or 3) outbreaks of other unidentified, and clinically milder, viruses within the genus *Ebolavirus*. The latter possibility was proposed by virtue of the fact that we only know of Taï Forest ebolavirus (TAFV) from a single case in humans ^16^, and that Bundibugyo ebolavirus (BDBV) was not discovered until 2007 ^17^, suggesting that our knowledge of the range of viruses in the genus *Ebolavirus* is far from complete. Recently, the number of known species in the genus *Ebolavirus* has expanded again with the discovery of Bombali ebolavirus (BOMV) ^18^. The wider family *Filoviridae* has also gained new members in Měnglà dianlovirus (MLAV) ^19^, Wenling frogfish filovirus ^20^ and *Rousettus leschenaultia* filovirus Bt-DH04 ^21^. It is therefore plausible that unknown filoviruses are circulating in Africa and Asia.

Explanation of the serological signal found in Germany in the early 1990s by Becker *et al* ^22^ is slightly more challenging. Although it is now established that EBOV may present occasional asymptomatic or very mild cases ^23–26^, it seems highly unlikely that entirely asymptomatic EBOV outbreaks could have occurred in Europe. If the results of Becker *et al* are produced by exposure within Europe to a filovirus, it cannot be EBOV or any of the known EVD-producing members of the family. There are now several known members of the family *Filoviridae* outside of Africa and, in Reston ebolavirus (RESTV) ^27^, an Asian member of the genus *Ebolavirus*, but the only filovirus known to be autochthonous to Europe is Lloviu cuevavirus (LLOV) ^28,29^ of bats. If there are no filoviruses circulating in Europe capable of infecting humans, an alternative explanation for the serological signal obtained by Becker *et al* (other than travel-related exposure) is an immunological cross-reaction between the filovirus antigens used in their study and antibodies against another virus to which their surveyed population had been exposed.

In this study, we attempt to replicate the findings of Becker *et al* using an anonymous set of sera donated in Lancaster, UK. We compare the results of our Lancaster volunteers with EBOV-produced EVD convalescent reference sera from the World Health Organization ^30,31^. PSI-BLAST ^32^ is used to identify homologues of EBOV nucleoprotein in non-filoviruses and structural alignment is performed. Finally, using PREDEP ^33^, we predict regions of potential antigenicity over stretches of structural homology.

## 2. Results

### 2.1 Calibration of ELISA using WHO Ebola reference sera

The EBOV vaccine ^34,35^ expresses EBOV glycoprotein within a recombinant VSV vector. Against experimentally presented EBOV glycoprotein antigen (GPZ), vaccinee serum has a stronger response on ELISA than either of the convalescent sera assayed (Figure 1; GPZ columns). As expected, vaccinee serum does not respond on ELISA to experimentally presented EBOV nucleoprotein antigen (NPZ), since nucleoprotein is not present in the vaccine (Figure 1; NPZ columns). By contrast, both convalescent sera samples generate an ELISA signal against both NPZ and GPZ, consistent with exposure to both glycoprotein and nucleoprotein during natural infection with EBOV. Furthermore, the strength of the convalescent sera response is three or more times higher for NPZ compared to GPZ, suggesting that the nucleoprotein is the more important antigen in the natural immune response to EBOV. Convalescent 2 was chosen as the positive control for further ELISAs based on its stronger signal against both antigens.

**Figure 1:**
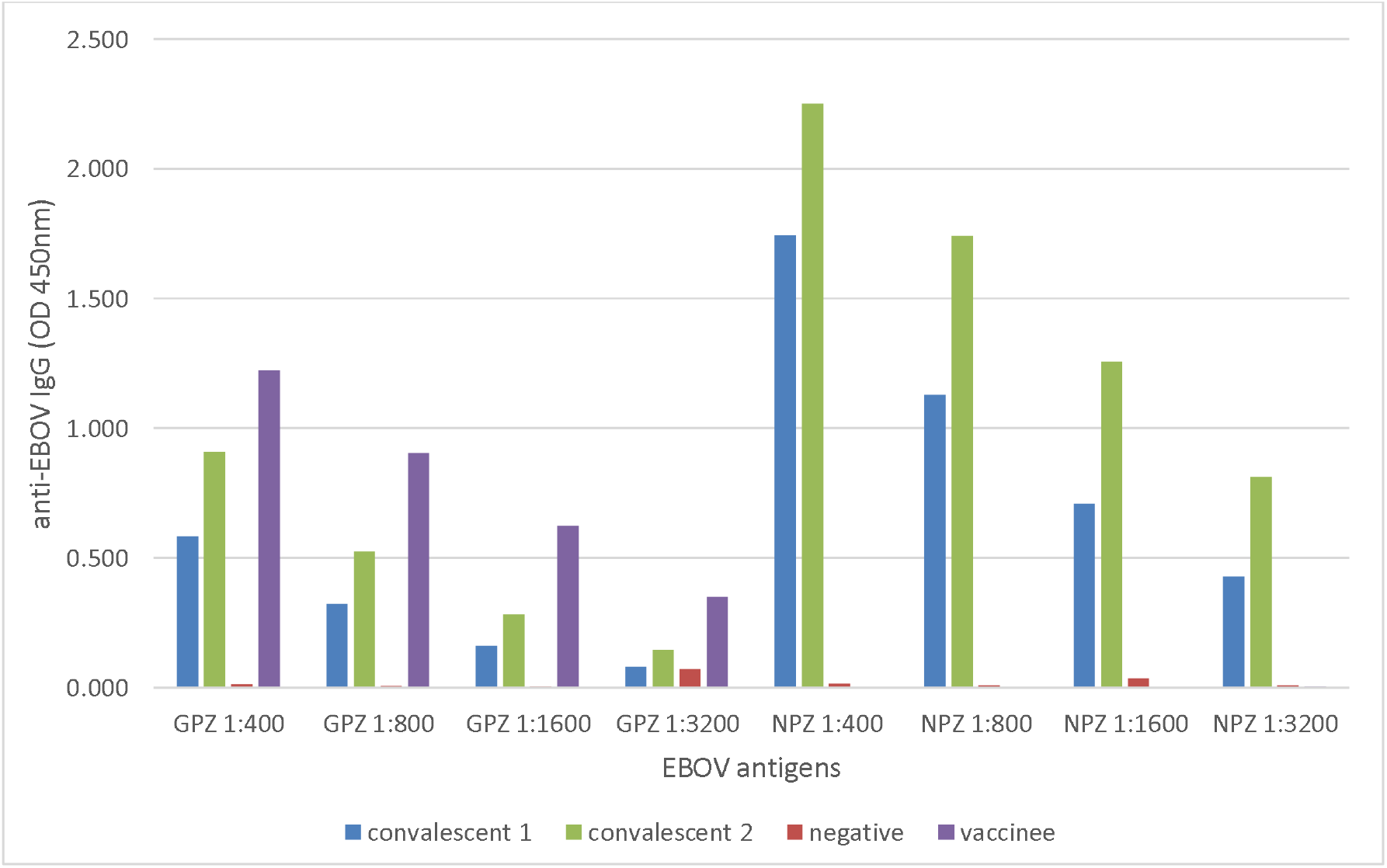
ELISA using known convalescent, vaccinee and negative control samples. GPZ: EBOV glycoprotein antigen; NPZ: EBOV nucleoprotein antigen.

### 2.2 ELISA against Lancaster-Morecambe donated sera

Since NPZ appears to elicit a stronger ELISA response in positive control sera (Figure 1), ELISA was then performed, using NPZ as the presented antigen, on our 144 Lancaster samples. Figure 2 shows the 25 samples, from the total 144, with the highest ELISA response to NPZ.

The three samples with the highest response to NPZ (Figure 2: P031, P028 and JP04) were then compared with the positive control convalescent sample (Figure 3). The convalescent positive control shows a stronger ELISA response to NPZ than the three Lancaster samples at all dilutions. The most positive Lancaster samples (P031) has ELISA OD_450_ levels ranging from two-thirds that of the positive control at dilution 1:200, to around one quarter at dilutions of 1:800 and below.

**Figure 2.**
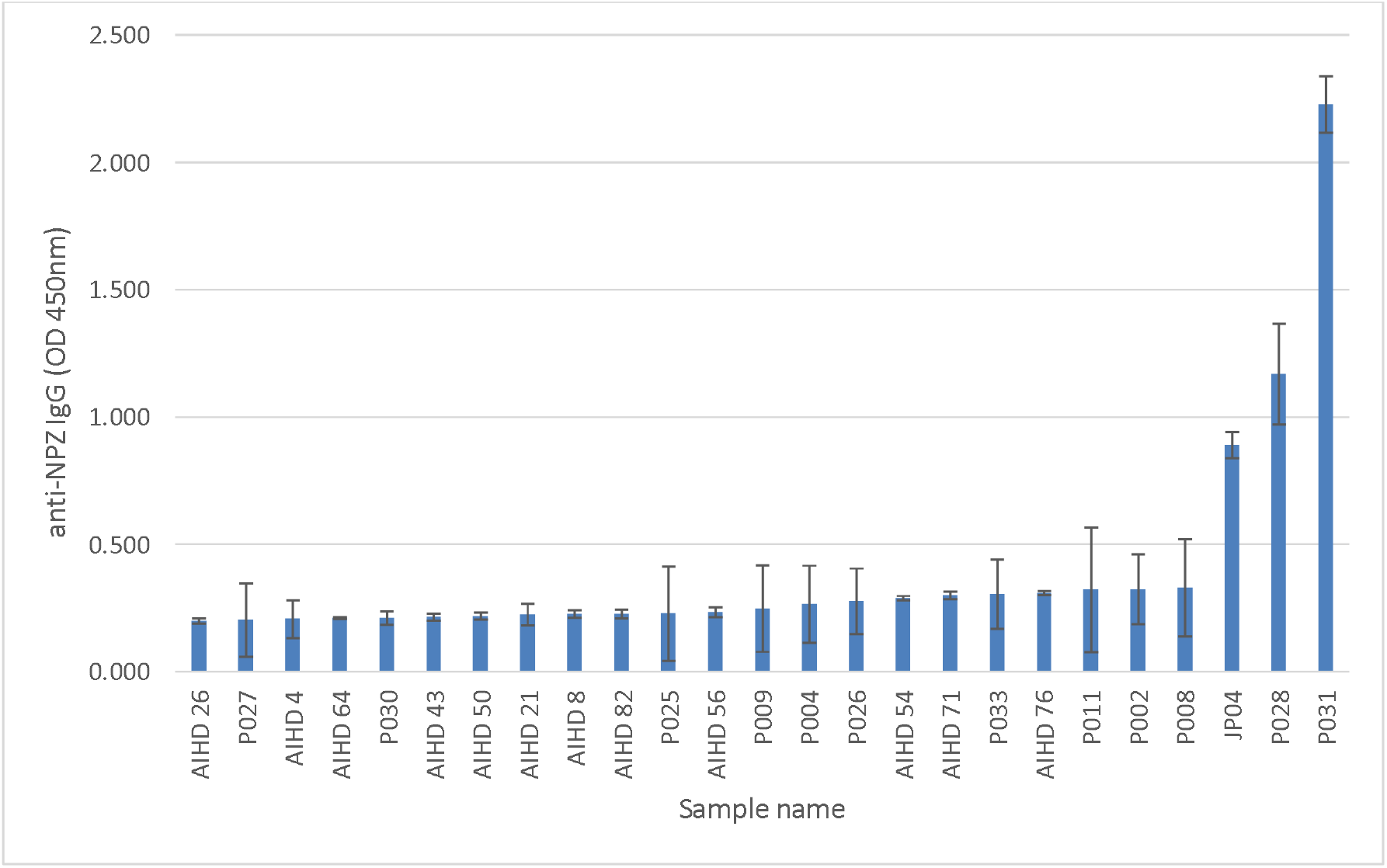
The 25 samples presenting with the strongest ELISA response to NPZ at 1:200 dilution. Error bars: standard deviation calculated from triplicate plating.

**Figure 3:**
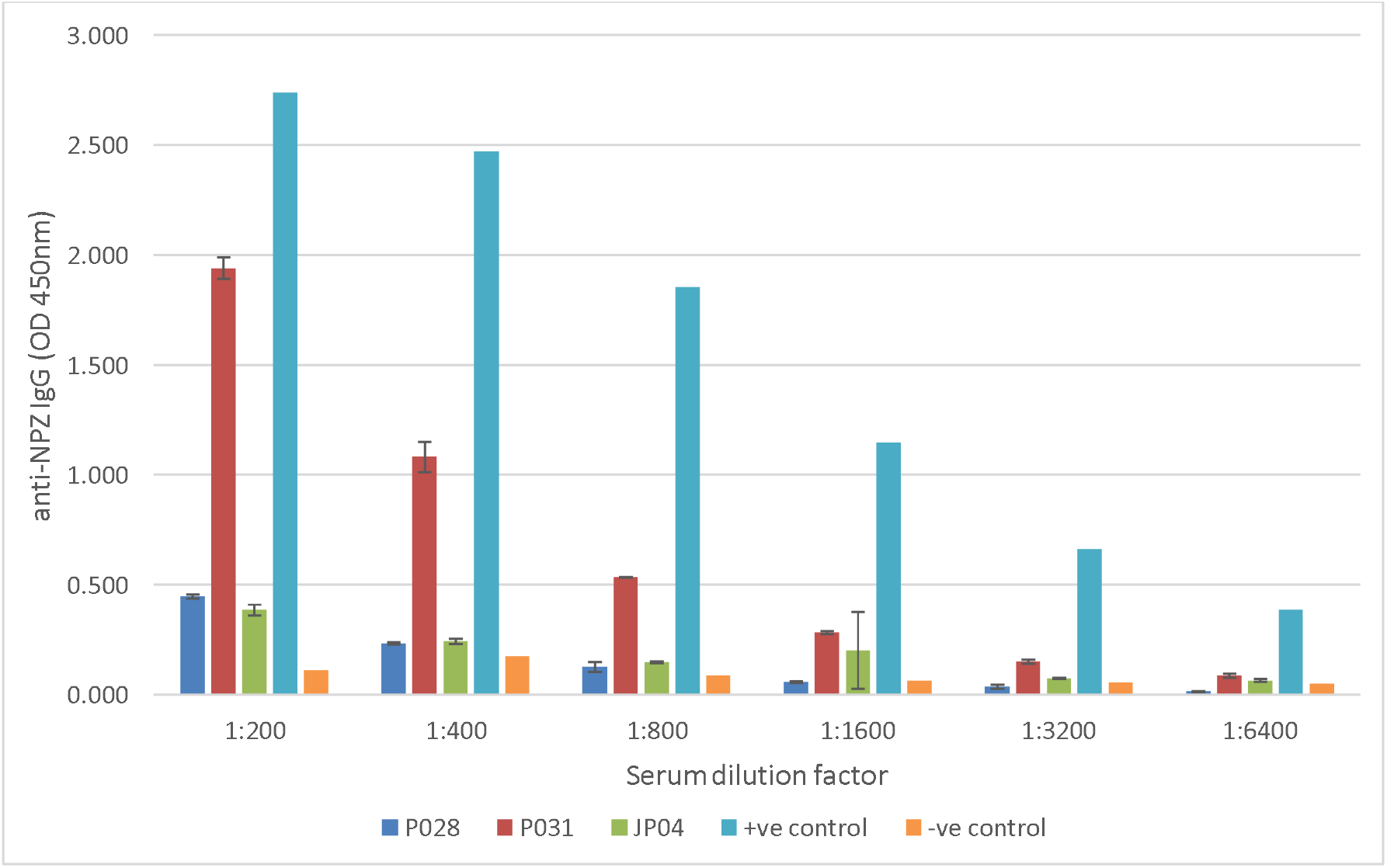
Comparison of Lancaster positive sera with convalescent positive control and negative control. Sera were tested against NPZ antigen. Error bars represent standard deviation calculated from triplicate plating. Positive control is convalescent 2 (see Figure 1).

The two samples showing the highest ELISA response to NPZ (Figure 2: P028 and P031) were tested against experimentally presented Sudan ebolavirus (SUDV) nucleoprotein (NPS) and SUDV glycoprotein (GPS) as well as NPZ and GPZ (Figure 4). Against the nucleoproteins (NPZ and NPS), P031 gives the strongest response, with ELISA OD_450_ levels nearly 4-fold higher than P028. By contrast, against the glycoproteins (GPZ and GPS) P028 is nearly 3-fold higher than P031. However, for both Lancaster sera the nucleoprotein antigens elicit the strongest response, as previously found for the convalescent sera (Figure 1).

**Figure 4:**
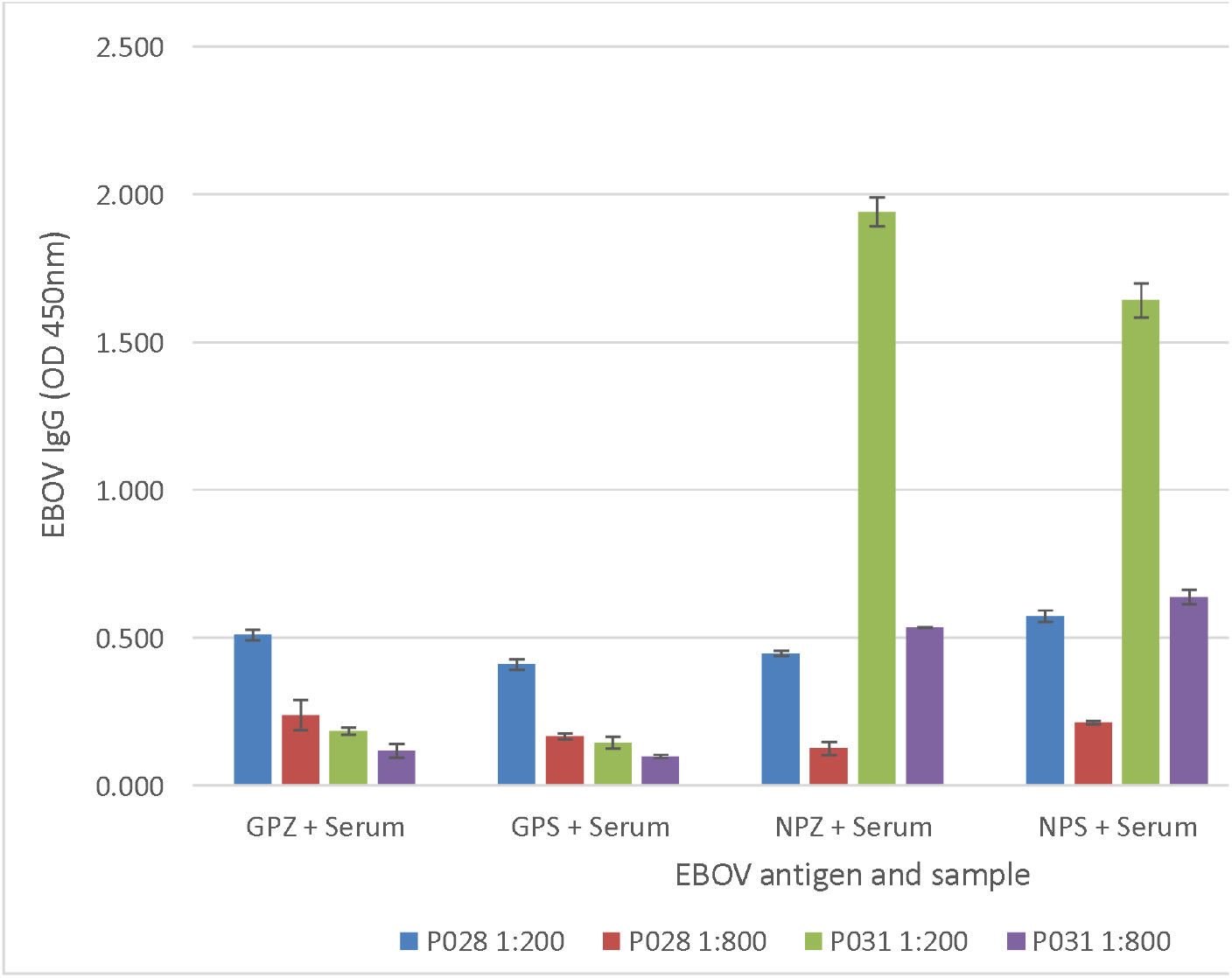
IgG profiles of donor samples P028 and P031. Each sample is tested against GPS, GPZ, NPS and NPZ and at two dilutions (1:200 and 1:800). GPS: Sudan ebolavirus glycoprotein; GPZ: Zaire ebolavirus glycoprotein; NPS: Sudan ebolavirus nucleoprotein; NPZ: Zaire ebolavirus nucleoprotein.

### 2.3 Western blot analysis of donor serum P031

Western blotting was performed using donor serum P031 and WHO reference positive control convalescent serum, and NPZ, GPZ, NPS and GPS antigens (Figure 5).

**Figure 5:**
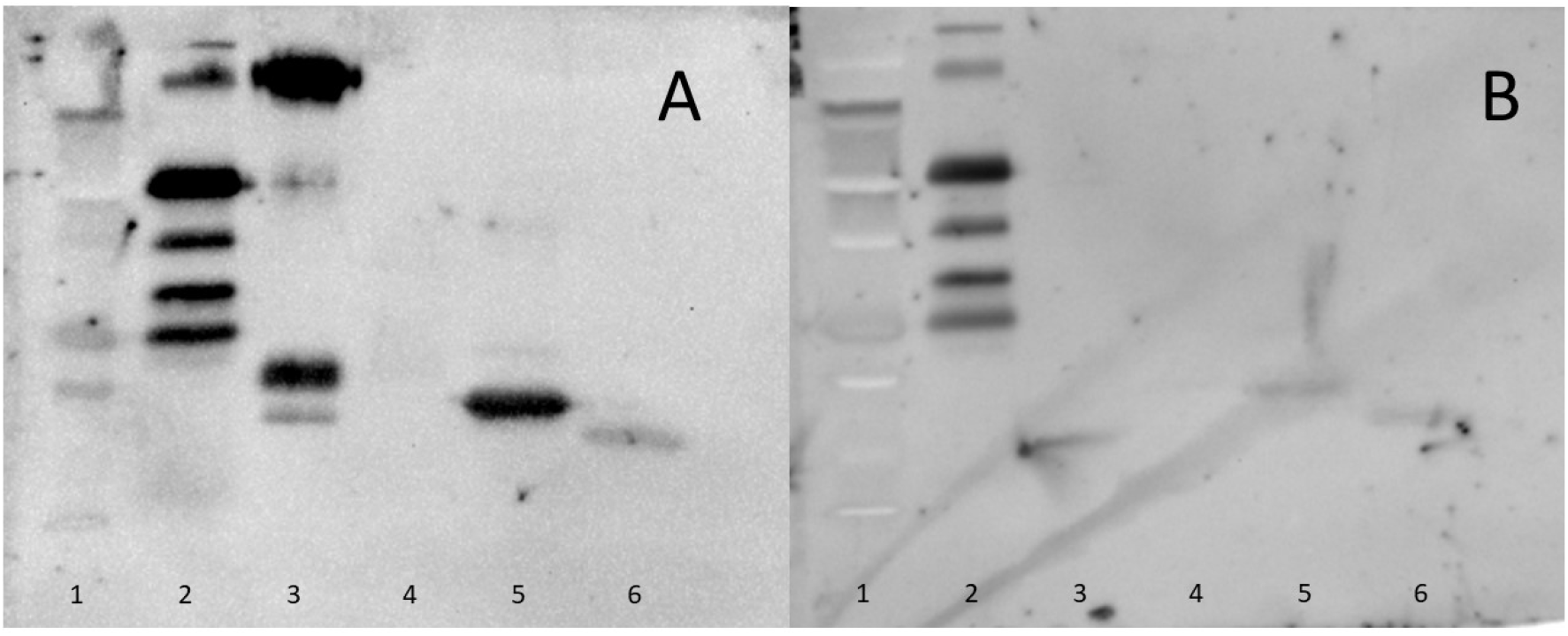
Western blot assay. A: WHO reference positive control convalescent serum. B: serum donor P031. Lanes as follows: 1: Low Molecular Weight Ladder, 2: IgG (positive control), 3: GPZ, 4: GPS, 5: NPZ, 6: NPS.

EBOV convalescent serum shows strong reactivity to GPZ (Figure 5A, lane 3) and NPZ (Figure 5A, lane 5). The response to GPS (Figure 5A, lane 4) appears negative, but there is a response to NPS (Figure 5A, lane 6). Donor serum P031 has no reactivity to GPZ or GPS (Figure 5B, lanes 3 & 4) but weak reactivity to NPZ and NPS (Figure 5B, lanes 5 & 6).

### 2.4 PSI-BLAST identifies closest non-filovirus relatives of EBOV nucleoprotein

PSI-BLAST indicates that the nearest relatives, in terms of amino acid sequence similarity, of filovirus nucleoproteins are the corresponding proteins of the family *Paramyxoviridae*, a sister family of *Filoviridae* within the order *Mononegavirales*. Measles morbillivirus and canine morbillivirus, the causative agents of measles in humans and distemper in dogs, respectively, have nucleoproteins with PSI-BLAST-generated local alignments displaying 15-18% sequence identity over 33-43% of the length of the nucleoprotein.

### 2.5 Structural similarities of EBOV nucleoprotein to other mononegaviral nucleoproteins

EBOV nucleoprotein has several solved crystallographic structures of which the longest is 4YPI ^36^, covering positions 38 to 385 of the 739 residue nucleoprotein. This N-terminal half of the nucleoprotein consists predominantly of alpha helices (Figure 6A). C-terminal to this point, the protein is predicted by GOR ^37,38^ to be mostly random coil with sporadic small alpha-helices and beta-strands. Superposition of EBOV nucleoprotein structure 4YPI with homologous solved structures from the order *Mononegavirales* shows that the closest overall structural similarity is with mammalian rubulavirus 5 (formerly parainfluenzavirus 5; PIV-5) nucleoprotein structure 4XJN ^39^ at 6.6 Å RMSD on structural superposition (Figure 6B and Table 1). However, plotting RMSD on a sitewise basis, using a sliding window of 25 residues, shows that measles morbillivirus nucleoprotein structure 5E4V ^40^ has a stretch commencing at position 242 and extending to position 354 in the alignment (equivalent to positions 245 to 353 in measles nucleoprotein), where the RMSD is always below 3.9 Å, and averages at 2.4 Å (Figure 6D). Even though the overall superposition of EBOV and measles morbillivirus nucleoprotein, at 14.5 Å, is poorer than that of EBOV and PIV-5 nucleoprotein (Table 1 and Figure 6B), the region spanning measles nucleoprotein positions 245 to 353 shows a visually striking superposition with EBOV nucleoprotein (Figure 6C).

**Table 1:** Root mean square deviation (RMSD) of mononegaviral nucleoprotein structures superposed with EBOV nucleoprotein. Values are in Ångstroms and calculated in MOE.

| Structure PDB id. | Family | Species | RMSD (Å) |
| --- | --- | --- | --- |
| 4XJN | *Paramyxoviridae* | Mammalian<br>rubulavirus 5<br>(formerly PIV-5) | 6.6 |
| 2GTT | *Rhabdoviridae* | Rabies lyssavirus | 10.3 |
| 2WJ8 | *Pneumoviridae* | Respiratory syncytial<br>virus (RSV) | 11.7 |
| 1N93 | *Bornaviridae* | Borna disease virus<br>(BDV) | 12.0 |
| 5E4V | *Paramyxoviridae* | Measles morbillivirus | 14.5 |

**Figure 6:**
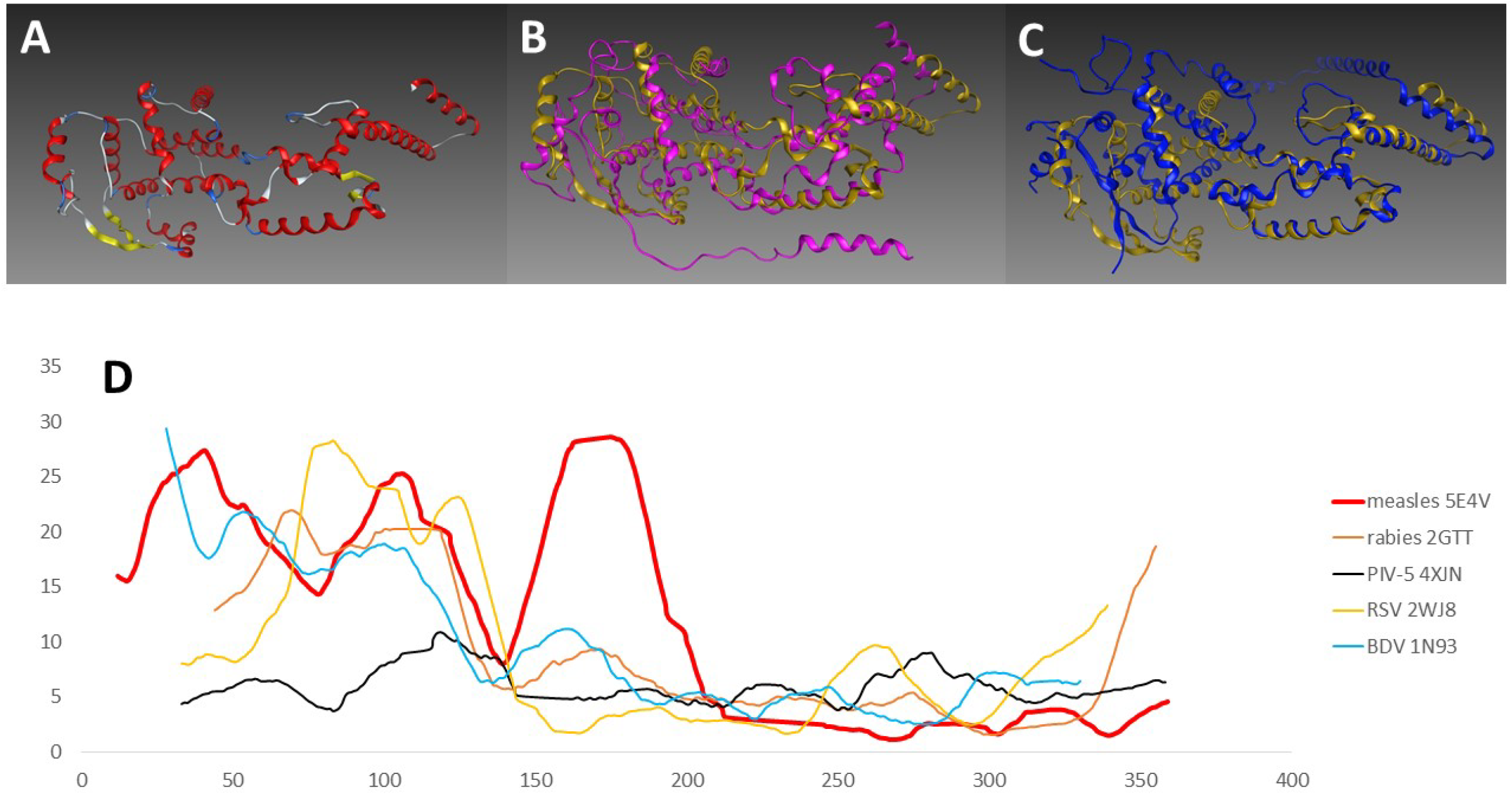
Structural superposition of mononegaviral nucleoproteins. A: The tertiary structure of EBOV nucleoprotein (PDB id. 4YPI.A) with alpha-helices in red and beta-sheets in yellow. B: EBOV nucleoprotein (4YPI.A, gold) superposed with parainfluenzavirus-5 nucleoprotein (4XJN.A, pink). C: EBOV nucleoprotein (4YPI.A, gold) superposed with measles morbillivirus nucleoprotein (5E4V.A, blue). D: Plot of the root mean square deviation (RMSD, sliding window length 25 residues) of the alpha-carbon atoms of the 5 mononegaviral nucleoprotein structures in Table 1 with EBOV nucleoprotein.

### 2.6 Antigenic epitope in region of closest structural superposition between EBOV and measles morbillivirus nucleoproteins

The region of highest structural similarity between EBOV and measles morbillivirus nucleoproteins has high potential to be an immune epitope. Table 2 summarises the best predicted epitopes produced when EBOV nucleoprotein is used as a query sequence for PREDEP. Two predicted epitopes binding to MHC alleles B3501-9 and A0201-10 overlap from positions 293 to 302, and two more binding to A0201-10 overlap from positions 307 to 324 (Table 2; Figure 7). EBOV and measles morbillivirus nucleoproteins are therefore predicted to possess epitopes with highly similar MHC specificities in a region of high structural similarity.

**Table 2:**
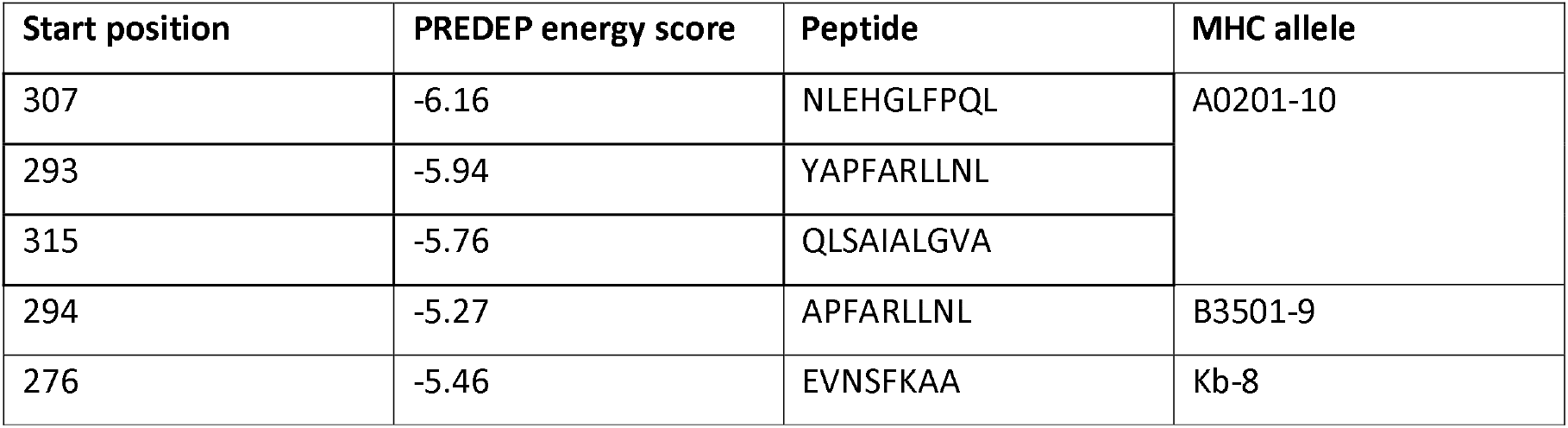
Best candidate epitopes within EBOV nucleoprotein identified by PREDEP. Start position in EBOV nucleoprotein amino acid sequence. See Figure 7 for localisation in superposed structures.

| Start position | PREDEP energy score | Peptide | MHC allele |
| --- | --- | --- | --- |
| 307 | -6.16 | NLEHGLFPQL | A0201-10 |
| 293 | -5.94 | YAPFARLLNL |  |
| 315 | -5.76 | QLSAIALGVA |  |
| 294 | -5.27 | APFARLLNL | B3501-9 |
| 276 | -5.46 | EVNSFKAA | Kb-8 |

**Figure 7:**
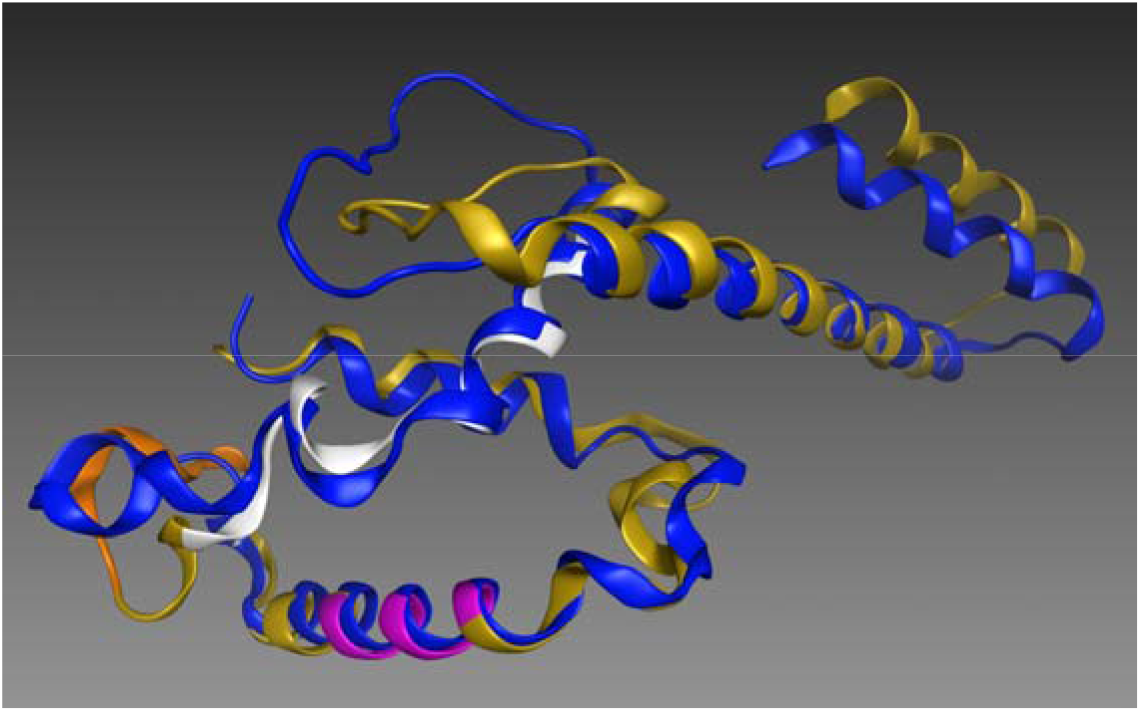
Antigenic epitopes identified by PREDEP. Detail of Figure 6C, showing only the region of maximum structural superposition between EBOV 4YPI (gold) and measles 5E4V (blue). PREDEP identified epitopes highlighted as follows: MHC allele Kb-8 (magenta); B3501-9 (orange); A0201-10 (white).

## 3. Discussion

Our ELISA experiments show strong signals from vaccinee sera for glycoprotein antigens and from convalescent sera for both glycoprotein and nucleoprotein antigens (Figure 1). This is consistent with the absence of nucleoprotein antigens in the recombinant vaccine. Within the convalescent sera, we see a stronger ELISA signal for nucleoprotein antigens. We believe that this may indicate that the nucleoprotein is the stronger antigen in the natural immune response to EBOV infection. However, we acknowledge that this may also be simply a reflection of the expression of the antigens in different culture systems (glycoproteins in HEK293 cells and nucleoproteins in *E.coli*), producing differing levels of antigenicity.

We also acknowledge that a weakness of our study is that it does not define precise thresholds for declaration that a sample is seropositive or seronegative for either antigen. Negative control sera show ELISA OD_450_ readings close to zero, and many of our Lancaster sera have responses four or five fold higher than negative controls (Figure 2). It might be tempting to postulate a high level of seropositivity, but we prefer to proceed cautiously and draw attention only to our three samples with signals comparable to positive controls. At a serum dilution of 1:200 against NPZ, these all have a mean ELISA OD_450_ reading greater than 0.75. Positive controls at serum dilution of 1:400 against NPZ have mean ELISA OD_450_ readings greater than 1.75 (Figure 1). Our strongest responder, patient P031, at serum dilution of 1:400 against NPZ has a mean ELISA OD_450_ reading of just over 1 (Figure 3). From the total 144 patients, we therefore propose that three (2.1%, 90% CI: 0.13%-4.07%) may be regarded as seropositive to NPZ. P031 is also positive to NPS and our second strongest responder to NPZ, patient P028, also shows seropositivity for GPS and GPZ (Figure 4).

Our confidence in our signals of seropositivity to nucleoproteins on ELISA is bolstered by the observation that a Western blot comparing P031 with a convalescent positive control shows bands of the correct size for the nucleoprotein (Figure 5, compare lanes 5 & 6 of A with 5 & 6 of B). By contrast, there is no evidence from our Western blots for any band corresponding to glycoprotein in P 031 (Figure 5, compare lanes 3 & 4 of A with 3 & 4 of B). The ELISA signal of glycoproteins is, however, weak in P031 compared to P028 (Figure 4), and we acknowledge that our study would have been strengthened by the inclusion of a Western blot comparing P031 with a convalescent positive control. Despite the shortcomings in terms of comprehensiveness of data, we believe that our analyses do indicate that we have discovered the first signal for seropositivity to filoviruses in Europe since the early 1990s. The incidence may conservatively be estimated at 2.1%, (90% CI: 0.13%-4.07%) and may have been estimated higher if we had been able to establish a definite threshold for seropositivity and thereby include more samples in the total (Figure 2). This seropositivity therefore requires an explanation.

The permanent resident population of the study area is 141,000 (45,000 urban, 96,000 rural), and is less ethnically diverse and more aged (95% white, 18% over 65 years) than the UK as a whole, or the north-west region of England. However, there are also approximately 20,000 temporary resident students, from two universities and various colleges, with a wide variety of origins. The possibility exists that some of the anonymous donor sera are from Africans, who may have previously encountered EBOV. Tourism and business travel to EVD risk regions of Africa from the UK is generally rare, but there is some family visit travel by UK residents of African origin. Additionally, a university population may include academics and students who have made equivalent field trips.

Even if some of our samples are from such individuals, the results obtained are insufficient to suggest exposure to EBOV. Although ELISA shows sample P028 to be positive for both filovirus nucleoproteins and glycoproteins (Figure 4), our Western blot on sample P031 does not confirm the ELISA glycoprotein signal, but only the nucleoprotein signal (Figure 5B). By contrast, EVD convalescent sera has a strong signal on Western blot to both glycoprotein and nucleoprotein antigens (Figure 5A). This suggests that a more probable explanation is that our Lancaster samples have been exposed to another virus which has generated antibodies capable of binding to EBOV, and to a lesser extent SUDV, nucleoprotein antigens.

It is possible that this hypothetical unknown virus is an undiscovered member of the family Filoviridae. Presumably this virus would have a mild clinical presentation, since anything approaching the kind of haemorrhagic fever symptoms seen in EVD would already have attracted considerable clinical attention. It would also need to be rare, since only 3 of 144 of our samples were assessed as positive in ELISA. Experimental isolation of such a virus might involve screening of sera from patients with fever who are negative for all microbiology and virology tests. Patients with respiratory symptoms might also be excluded on the grounds that their fever is less likely to be caused by a virus within the family Filoviridae. For a common virus, the target study population for virus isolation and sequencing would be infants, as older patients would be more likely to be already immune. However, since we are suggesting a virus with approximately a 2% population seropositivity, presumably reflecting a 2% lifetime incidence, early exposure might be uncommon. In this scenario, a more even spread across age groups might be expected, thus making identification of a target study population more difficult.

Alternatively, the virus generating the antibody response may not be a member of the family Filoviridae. It has been known for several years that nucleoproteins exhibit sequence and structural similarity across distinct virus families ^41,42^. Our bioinformatics analysis suggests that the region of this structural similarity covers a potential epitope region. Antibodies generated against these epitopes may thus be posited to cross-react with the nucleoproteins of filoviruses. Measles morbillivirus would appear to be a particularly likely candidate to satisfy this theory as its nucleoprotein is the closest relative among the paramyxoviruses to the nucleoprotein of EBOV, as assessed by PSI-BLAST, and possesses a region of striking structural similarity in a candidate epitope region (Figure 6 & 7; Table 2). The only clinical details available for our three candidate positive samples are that cases JP04 and P031 both had pneumonia and P028 a caecal volvulus. Although pneumonia is a known complication of measles, the characteristic rash would surely have been noted and a more precise diagnosis obtained for these two patients. The cross-reactivity must therefore stem from an earlier encounter. The incidence of exposure to measles antigens, which primarily occurs in the UK through MMR vaccination together with a residue of live cases in unvaccinated individuals, approaches 90%. The theory of anti-measles morbillivirus nucleoprotein antibody cross-reactivity with EBOV nucleoprotein antigens therefore begs the question of why the incidence of such cross-reactivity is only just over 2%. It is known that infection with live measles morbillivirus in non-immune individuals elicits a stronger antibody response than those receiving the attenuated strain in the MMR vaccine ^43^. In Africa, measles outbreaks are still common, especially when vaccination programmes fail due to civil conflict or overstretching of medical systems during other epidemics, as happened in the aftermath of the EVD outbreak in west Africa ^44–46^. However in England, confirmed cases of measles are only in the order of several hundred per year with two-thirds of cases in the over 15 age group ^47^, so recent live measles infection cannot account for the 2% seropositivity to EBOV nucleoprotein observed here. It is therefore necessary to posit that cross-reactivity perhaps originates in childhood MMR vaccination and gradually wanes over several years, producing a low incidence in adults. This is a testable hypothesis, which could be confirmed by detecting a higher incidence of seropositivity to EBOV antigens in recently vaccinated children than in adults who received MMR two decades ago.

## 4. Methods

### 4.1 Serum collection

144 anonymised human serum samples were collected in the Lancaster and Morecambe area (54°N, 2.8°W), from intensive care unit (ICU) patients of local hospitals, during 2012 and 2013. 100 were collected from ischaemic heart disease patients (AIHD 1-100), and the remaining 44 samples (P001-039 and JP01-05) were obtained from patients with a variety of pathologies. Gender, ethnic origins of donors and their travel history were not recorded. After collection, the samples were kept at −80°C.

### 4.2 Antigens

ELISA and Western blots used recombinant EBOV glycoprotein (GPZ, 2B Scientific, IT-014-003p), recombinant SUDV glycoprotein (GPS - Nakisamata, 2B Scientific, IT-014-007p), recombinant EBOV nucleoprotein (NPZ, AMS Bio, R01578-1) and recombinant SUDV nucleoprotein (NPS, AMS Bio, R01577-1). These manufacturers expressed the glycoproteins in human embryonic kidney cell line (HEK293) culture, and the nucleoproteins in *E.coli*.

### 4.3 Antibodies and control sera

For the detection of primary antibodies in serum samples, a polyclonal, HRP-conjugated goat anti-human-IgG secondary antibody was used (Abcam, ab6858). Convalescent serum from a patient in the 2013-2016 EVD outbreak in West Africa, double detergent extracted and certified as virus-free by the National Institute of Biological Standards and Controls (NIBSC) UK, was used as positive control ^30,31^. The negative control was commercial pooled human serum (Sigma-Aldrich).

### 4.4 Enzyme-linked Immunosorbent Assay (ELISA)

Indirect ELISA ^48^ was used to detect anti-EBOV IgG in human serum samples, as follows. 96-well microplates (Nunc Maxisorp) were coated with antigen (0.5 μg/mL) in Carbonate Coating Buffer (Sigma Aldrich) overnight at 4°C. The plates were then washed three times with phosphate-buffered saline solution with 0.05% Tween 20 (PBST). To block non-specific binding to antigens, the wells were then treated with 200 μL/well of 5% w/v milk powder in PBST, at 37°C for 1 hour, followed by three further washes with PBST. Sera, positive and negative controls were diluted in PBST. 100 μL/well was plated and incubated at 37°C for 1 hour, then wells washed again as above. The secondary antibody was added at 1:4000 in PBST, and incubated again at 37°C for 1 hour, followed by a further washing step and the addition of the SureBlue™ (TMB). After 7 minutes, the reaction was stopped by the addition of 1N sulphuric acid. The optical density (OD) was determined at 450nm for 0.1 seconds using a Wellac Victor spectrophotometer.

### 4.5 Western blot

Antigens were diluted to 5 μg/mL and heated for 3 minutes at 98°C. 20 μL of each antigen solution was added per lane of a polyacrylamide gel, and run at 140V for 60 minutes. Immunoblot membranes were equilibrated by soaking in methanol for 20 seconds, followed by 2 minutes in H_2_O and finally a minimum of 20 minutes in transfer buffer (45 g Glycine, 9.7 g Tris, 800mL methanol, made to 4L with dH_2_O). The transfer stack was soaked in transfer buffer and gels were transferred at 115V for 60 minutes. Membranes were then blocked with 5% w/v milk solution for 1 hour at room temperature on an orbital shaker and washed with PBST. The serum samples were then diluted in PBST at 1:1000. 10 mL was added to each membrane and shaken overnight at 4°C, followed by a 30-minute washing of the membranes in PBST. The secondary antibody (Ab6858) was added as a 1:4000 dilution in PBST and shaken at room temperature for 60 minutes. Membranes were washed 3 times for 15 minutes with PBST followed by the addition of the detection reagent (Pierce ECL WB Substrate) and development (ChemiDoc XRS+, Bio-Rad).

### 4.6 Bioinformatics

Identification of non-filovirus nucleoproteins with sequence similarity to EBOV nucleoprotein was performed using PSI-BLAST ^32^, as follows: BLASTP was run in PSI-BLAST mode with a maximum of 100 hits, using EBOV nucleoprotein (accession NP066243.1) as a query, and was configured to exclude hits from EBOV. From the results of the first PSI-BLAST round, one nucleoprotein sequence was selected each from other members of the genus Ebolavirus (AGL73450.1, YP003815423.1, AGC02891.1, YP009513274.1, ACR33187.1 and YP004928135.1), and a PSI-BLAST second iteration run, again with 100 hits maximum. Hits below the threshold of the first iteration now appear, of which the top hit is ALL29056.1 – the nucleoprotein of measles virus genotype B3.

Alignment and structural superposition of viral nucleoproteins with solved X-ray crystallographic structures was performed in Molecular Operating Environment (MOE), 2018.01 (Chemical Computing Group ULC, 1010 Sherbooke St. West, Suite #910, Montreal, QC, Canada, H3A 2R7). Antigenic epitopes were identified using PREDEP ^33^.

## 5. Acknowledgements

This work was funded by a Lancaster University Early Stage Career Grant (ESCG) to DG. We thank visiting intern Katharina Hartman (Jacobs University Bremen, Germany) and undergraduate Magdalena Formella (Lancaster University) for laboratory assistance in the preliminary stages of this project, and Dianna Wilkinson and Stacey Efstathiou (NIBSC, Ridge, Herts., UK) for provision of WHO convalescent positive control reference sera. Thomas R.W. Tipton and Miles W. Carroll (Public Health England, Salisbury, UK) provided many useful technical comments on the manuscript and general discussion of the meaning of the results.

## 6. Author Contributions

LAB performed and supervised laboratory work. MM performed laboratory work. AB-K performed bioinformatics. RML was executive supervisor for laboratory work, co-supervisor on the MSci project of MM and named individual on IRAS 117608. DG wrote the manuscript, performed bioinformatics, was co-supervisor on the MSci project of MM and supervisor on the BSc project of AB-K.

## 7. Ethical Statement

The sample collection protocol was approved by the Research Ethics Service of the NHS in England (IRAS application number 117608). The original project from which donated samples were obtained, was funded by University Hospitals of Morecambe Bay NHS Foundation Trust.

